# CaptureSeq: Hybridization-based enrichment of *cpn60* gene fragments reveals the community structures of synthetic and natural microbial ecosystems

**DOI:** 10.1101/492116

**Authors:** Matthew G. Links, Tim J. Dumonceaux, Luke McCarthy, Sean M. Hemmingsen, Edward Topp, Jennifer R. Town

**Author notes:** Corresponding Author: Jennifer R. Town, 107 Science Place, Saskatoon, Saskatchewan S7N 0X2 Canada.

## Abstract

**Background:** Molecular profiling of complex microbial communities has become the basis for examining the relationship between the microbiome composition, structure and metabolic functions of those communities. Microbial community structure can be partially assessed with universal PCR targeting taxonomic or functional gene markers. Increasingly, shotgun metagenomic DNA sequencing is providing more quantitative insight into microbiomes. However, both amplicon-based and shotgun sequencing approaches have shortcomings that limit the ability to study microbiome dynamics.

**Methods:** We present a novel, amplicon-free, hybridization-based method (CaptureSeq) for profiling complex microbial communities using probes based on the chaperonin-60 gene. Molecular profiles of a commercially available synthetic microbial community standard were compared using CaptureSeq, whole metagenome sequencing, and 16S universal target amplification. Profiles were also generated for natural ecosystems including antibiotic-amended soils, manure storage tanks, and an agricultural reservoir.

**Results:** The CaptureSeq method generated a microbial profile that encompassed all of the bacteria and eukaryotes in the panel with greater reproducibility and more accurate representation of high G/C content microorganisms compared to 16S amplification. In the natural ecosystems, CaptureSeq provided a much greater depth of coverage and sensitivity of detection compared to shotgun sequencing without prior selection. The resulting community profiles provided quantitatively reliable information about all three Domains of life (Bacteria, Archaea, and Eukarya) in the different ecosystems. The applications of CaptureSeq will facilitate accurate studies of host-microbiome interactions for environmental, crop, animal and human health.

## Introduction

Life on Earth is classified into hierarchical taxonomic lineages that describe all living systems as having descended from a common ancestor along three evolutionary lines. Using ribosomal RNA-encoding gene sequences, Woese & Fox (1977) delineated these Domains, which are now known as Bacteria, Archaea, and Eukarya (Woese et al. 1990). Most complex microbial communities exist as assemblages replete with representatives from each of these Domains, the total genomic complement of which is called a microbiome. Understanding microbial community dynamics requires tools to examine the composition of these complex ecosystems. Advancements in DNA sequencing technology have created new opportunities to simplify the profiling of microbial communities from a diverse range of environments. Insights gained through the study of the diversity of microbiomes in soil, water, plant and animal-associated ecosystems have revealed the powerful effects that microbiome composition and structure can have on how these communities function (Tikhonovich & Provorov 2011). A comprehensive understanding of the multifaceted relationships between microorganisms and their environment requires the generation of microbial community profiles that reflect as accurately as possible the original composition and quantitative structure of the microbial ecosystem under analysis.

In adapting the use of PCR for amplifying a conserved region of 16S rRNA, Weller and Ward provided the first example of microbial profiling (Weller & Ward 1989). Since then, microbiologists have increasingly embraced such culture-independent methods of identification (J T Staley & Konopka 1985). PCR-based amplification of 16S rRNA-encoding genes has become the method of choice for determining the composition of bacterial communities in a wide range of ecological niches (Edge et al. 2020; Turnbaugh et al. 2007). More recently, Paul Hebert’s proposed DNA barcoding criteria for Eukarya have established standards for what comprises a robust target for phylogenetic profiling (Hebert et al. 2003). Alternative universal gene markers for 16S (Singer et al. 2016), *cpn60* (Hill et al. 2004), *rpoB* (Adékambi et al. 2009), *mcrA* (Barret et al. 2013) and ITS (Schoch et al. 2012) have been used for profiling microorganisms from bacterial, archaeal and eukaryotic Domains, although no single amplification is able to profile microbes from all three Domains simultaneously. In order to obtain phylogenetic information for microorganisms across all three Domains of life, separate target amplification and processing protocols are required (Barret et al. 2015), increasing the cost and analytical complexity of accurately assessing dynamic changes in the community across Domains. Moreover, stochastic effects of primer interaction with a complex template, along with the difficulty in designing primers and amplification conditions that will equally target all members of a community (Walker et al. 2015), result in an unavoidable bias in community representation both in terms of presence/absence and relative abundances (Guo et al. 2016; Lynch & Neufeld 2015; Poretsky et al. 2014; Walker et al. 2015).

In recent years, metagenomic approaches in which whole nucleic acid recovered from a sample is fragmented and sequenced using “shotgun” methods have become increasingly popular. This approach has a significant advantage over methods based on amplification of taxonomic markers in that shotgun-sequencing data can overcome issues of bias and representation that are inherent in amplicon sequencing approaches, and it provides the additional advantage of describing the metabolic potential of the microbial community (Handley et al. 2013; Hess et al. 2011; Raymond et al. 2016). Sequencing of all DNA present in an environmental sample can therefore be considered a “gold standard” for taxonomic profiling. However, this approach is not without its own limitations. For example, it can be a wasteful enterprise in terms of the phylogenetic information recovered per sequencing cost. Shotgun sequencing is also not easily able to connect the functional potential observed in the sequencing data with the exact microbe within which that functionality resides. Additionally, DNA acquired from a community of microorganisms is inherently unbalanced; there are not equal numbers of each taxon, nor do all taxa have genomes that are of equal sizes. Thus shotgun sequencing can provide a view of microbial community composition that is biased by genome size and microbial abundances. Overcoming this bias requires significant amounts of sequencing; therefore, chasing the rarity of the least abundant microbes by shotgun metagenomics sequencing carries a high financial cost (Fierer et al. 2012; Guo et al. 2016; Luo et al. 2014; Lynch & Neufeld 2015). The abundances of microbes within characterized complex microbial communities range over many orders of magnitude. While shotgun sequencing efforts provide a reasonable estimate of abundance there is a significant loss in dynamic range when compared to PCR-based profiling.

The chaperonin 60 gene (*cpn60*) (Hill et al. 2004) (type I chaperonin) and its Archaeal homologue thermosome complex (Chaban & Hill 2012) (type II chaperonin) have been previously recognized as highly discriminating targets across all Domains of life (Links et al. 2014), meet standard International Barcode of Life criteria (Links et al. 2012), and enable the *de novo* assembly of operational taxonomic units (OTU) (Links et al. 2013). While “universal” PCR primers are available (Hill et al. 2004; Hill et al. 2006), they are not expected to capture the pan-Domain diversity of a complex microbial community through amplification. Moreover, *cpn60* amplification provides OTU abundances that do not always correlate to the true abundance of the microorganism in the sample (Dumonceaux et al. 2006). If these limitations can be overcome, there is significant opportunity to improve dramatically research assessing host-microbiome interactions in plant, human and animal settings.

Recent advances in hybridization-based DNA capture combined with high throughput sequencing (CaptureSeq), which have proven to be remarkably powerful means of enriching samples for DNA sequences of interest (Gasc & Peyret 2018; Schuenemann et al. 2011; Wagner et al. 2014), led us to consider the possibility of exploiting the unique features of *cpn60* to provide a microbial community profile without the use of universal PCR amplifications. A custom array of biotinylated RNA capture baits was designed based on the entire taxonomic composition of the chaperonin database cpnDB (www.cpndb.ca) (Hill et al. 2004; Hill & Vancuren 2019) and evaluated as a tool for enriching total genomic DNA simultaneously for type I and type II chaperonin target sequences. The features of the CaptureSeq method were determined in relation to results obtained using shotgun metagenomic sequencing and amplification of 16S rRNA-encoding genes on synthetic and natural microbial communities spanning a range of microbial ecosystems. Moreover, CaptureSeq was used to profile soil samples that have been treated with antibiotics over a 15 year time period. The results indicate that CaptureSeq provides the taxonomic reach associated with shotgun metagenomic sequencing combined with the sampling depth of amplicon-based sequencing.

## Materials & Methods

### CaptureSeq array design

Capture probes were designed based on all type I and type II chaperonin sequences in the public domain (i.e. cpnDB; www.cpndb.ca) (Hill et al. 2004). 15,733 probes were designed to be complementary to the type I and type II chaperone sequences. Design of probes was based on identifying 120 bp sequences from the reference database using a 60 bp incrementing step. Thus the resulting probes share a 50% overlap with the next probe in a tiling-like fashion. The custom biotinylated RNA oligonucleotides were provided in equimolar concentrations as a pooled Mybaits array by Arbor Biosciences (Ann Arbor, MI, USA).

### Template DNA preparation

The Zymobiomics microbial community DNA standard consisting of 8 bacterial and 2 fungal genomic DNAs (cat. no. D6305) was obtained from Zymo Research (Irvine, CA). This standard was diluted 1:20 as recommended by the manufacturer for 16S-based amplicon analysis and was also prepared for whole metagenome sequencing and CaptureSeq as described below.

A second synthetic microbial community was prepared using *cpn60* plasmids spiked into a naturally occurring microbial ecosystem. Background genomic DNA was prepared by washing wheat seeds and extracting DNA as previously described (Links et al. 2014). Amplicons corresponding to the *cpn60* universal target (*cpn60* UT) of 20 bacteria associated with the human vaginal tract (Links et al. 2013) and known to be absent from the seed wash DNA background (Links et al. 2014) were cloned into the pGEM-T Easy plasmid (Promega, WI, USA) and purified using the Qiagen Miniprep kit (Qiagen, CA, USA). The synthetic community was formed by combining equimolar concentrations of plasmids containing the *cpn60* UT for all 20 microorganisms (Links et al. 2013). Dilutions of this mixture (corresponding to 0.4, 0.04, and 0.004 ng plasmid DNA, or approximately 10^8^, 10^7^, and 10^6^ copies of each plasmid) were spiked into a background of 10 ng/μl of wheat seed carrier DNA. Spiked genomic DNA samples prepared in this way were sequenced using *cpn60* universal target amplification and CaptureSeq as described below.

Soil samples were obtained from a long-term study initiated in 1999 evaluating the effect of annual antibiotic exposure on soil microbial communities, described in Cleary et al. (2016). Soil samples evaluated in the present study were obtained in 2013 following 15 sequential annual applications of a mixture of sulfamethazine, chlortetracycline and tylosin, each added at concentrations of 0.1, 1, or 10 mg kg^−1^ soil, along with untreated control plots. Soil was sampled 30 days after the spring application of antibiotics. The plots were planted with soybeans (*Glycine max*, v. Harosoy) immediately after incorporation of the antibiotics. Each treatment level was applied to triplicate plots yearly since 1999 as described (Cleary et al. 2016). Genomic DNA was extracted from 3.5 g of each soil sample using the PowerMax Soil DNA isolation kit (Mo-Bio Laboratories, Carlsbad, CA) with a 5 mL elution volume. DNA extracts were quantified using a Qubit fluorimeter (Thermo Fisher Scientific, Waltham, MA, USA) and stored at −80°C until processing and analysis.

A water sample was obtained from a pond located on a Saskatchewan farm (51.99°N, - 106.46°W) on May 13, 2016. Biological material was recovered from 2L of water by centrifugation at 20,000 *g* for 20 minutes. Total DNA was extracted using a PowerWater DNA extraction kit (Mo-Bio Laboratories, Carlsbad, CA) and quantified as described above.

Samples were obtained from bovine manure storage tanks after 28 weeks of storage, as part of a separate study examining the effects of storage parameters on the methanogenic communities. DNA was extracted from 1 ml of the slurry using a commercial kit (Qiagen).

### Amplicon-based microbial community profiling

16S rRNA-encoding genes were amplified using primers 515f/926r under the recommended conditions (Caporaso et al. 2018). Amplicons were processed for sequencing using the NEBNext Illumina library preparation kit (New England Biolabs, Whitby, ON, Canada) and sequenced with 2×250 bp cycles of v2 Miseq chemistry (Illumina, San Diego, CA). Replicate reactions from each sample were pooled and gel purified using the Blue Pippin Prep system (Sage Science, MA, USA) with a 2% agarose cassette, and concentrated using Amicon 30K 0.5 ml spin columns (EMD Millipore, MA, USA). 16S amplicon from all samples was prepared for sequencing using the NEBNext Illumina library preparation kit (New England Biolabs), and sequenced with 400 forward cycles of v2 Miseq chemistry.

### Whole metagenome and CaptureSeq sample preparation

Genomic DNA was diluted to 2.5 ng/μl and split into two aliquots of 100 μl each for shearing using a water bath sonicator as described (Dumonceaux et al. 2017). Shotgun metagenomic sequencing libraries were prepared directly from one aliquot of each sheared genomic DNA sample using the NEBNext Illumina library preparation kit according to the manufacturer’s directions (New England Biolabs, MA, USA). Samples were then sequenced with 2×250 bp cycles of v2 Miseq chemistry (Illumina, CA, USA).

To generate the CaptureSeq libraries, the second aliquots of sheared genomic DNA samples were subjected to end repair and index addition using NEBNext as above, then hybridized to the capture probe array as described (Dumonceaux et al. 2017). The chaperonin-enriched products were then sequenced with 2×250 bp cycles of v2 Miseq chemistry (Illumina, CA, USA).

### Reference mapping

A reference database of all publicly available chaperonin sequences was generated by selecting a list of seven chaperonin protein sequences representing each taxonomic group: fungi, bacteria, archaea, plant mitochondria, plant chloroplast, and animal mitochondria. These sequences were used as queries for a BLAST search of GenBank using the default parameters to blastp. Matching protein sequences were manually vetted to generate a list of 30,141 protein identifiers. These protein identifiers were then used to retrieve the corresponding 30,120 nucleotide sequences available in GenBank according to the procedure described in Supplemental file S1. The accession numbers of those nucleotide sequences are provided in Supplemental Dataset S1. The breadth of taxa that were retrieved by this method was similar to the taxonomic breadth represented in the 16S and ITS reference datasets (Supplemental Dataset S2). Sequencing reads from all samples were grouped into taxonomic clusters by paired local alignment to this reference set of chaperonin genes using bowtie2 (v. 2.2.3) (Langmead et al. 2009). The sequencing libraries were down-sampled to the size of the smallest shotgun metagenomic library (2,777 mapped paired reads), and the number of reads mapping to each of the resulting taxonomic clusters was used as the basis for assessing the alpha and beta diversity metrics of the two profiling methods for equivalent sampling effort.

To compare the number of output sequencing reads for the different spiking levels, sequencing reads from the synthetic community-spiked samples were down-sampled to the smallest library size for each profiling technique (30,091 for amplicon and 506,247 for CaptureSeq) and mapped to a reference set of *cpn60* UT sequences for the 20 microorganisms in the panel by local paired alignment using bowtie2 as above.

### Sequence assembly

Read pairs from target taxonomic clusters obtained by reference mapping as described above were assembled *de novo* into *cpn60* OTU using Trinity (v. 2.4.0) with a kmer of 31 as described (Links et al. 2013).

### Sub-OTU (sOTU) definition using amplicon sequence variants (ASV)

For 16S sequencing reads, primer sequences were trimmed using cutadapt (v2.8) (Martin 2011), merged using FLASH2 (v2.00) (Magoc & Salzberg 2011), and ASV were determined using DADA2 (Callahan et al. 2016). For CaptureSeq reads, the *cpn60* sOTU was defined as nucleotides 1-220 of the *cpn60* UT (after trimming the 5’ amplification primer). Primer sequences were removed and all reads were trimmed to 220 bp using cutadapt. Sequences in the reverse orientation were reverse complemented prior to ASV analysis with DADA2.

### Alpha diversity analysis

To compare the richness and diversity metrics between the three profiling techniques, mapped sequencing reads were down-sampled from 250-2,750 reads to simulate a uniform sampling effort across profiling techniques. Metrics were averaged across 100 bootstrapped datasets using the multiple_rarefactions.py and alpha_diversity.py scripts from QIIME (v. 1.8.0) (Caporaso et al. 2010). Statistical significance between alpha diversity metrics was determined using the Kruskal-Wallis rank-sum test and equality of variances was evaluated using Lavene’s test from the stats and rstatix packages in R.

In the cases where the total effect of sequencing effort was required for comparisons across estimates of community coverage, read thresholds were transformed to reflect total sequencing effort for each sample.

### Beta diversity analysis

To compare the community similarity between different sequencing methods, mapped sequencing reads were down-sampled to the size of the smallest metagenomic library sample (2,777 mapped reads). For intra-technique comparisons, mapped sequencing reads were down-sampled to the smallest library size within each profiling method; 2,777 for metagenomic, and 127,642 for CaptureSeq. Principal Coordinate Analysis of inter- and intra-technique Bray-Curtis distance was calculated using the vegan package (v. 2.4.2) in R (v. 3.2.4).

### OTU quantification

Assembled OTU-specific primer and hydrolysis probe sets were designed using Primer3 (Rozen & Skaletsky 2000) or Beacon Designer (v.7) (Premier Biosoft, Palo Alto, CA, USA) as described previously (Pérez-López et al. 2017). Annealing temperatures were optimized for each reaction using gradient PCR with ddPCR Supermix for Probes (Bio-Rad, Mississauga, ON, Canada) using 900 nM each primer and 250 nM of hydrolysis probe in a 20 μl reaction volume. Primer/probe sequences and optimized amplification conditions are shown in Table S1. Template DNA was digested prior to amplification using *Eco*RI at 37°C for 60 minutes. A final volume of 2-5 μl was used as template for droplet digital PCR (ddPCR). Emulsions were formed using a QX100 droplet generator (Bio-Rad, Hercules, CA, USA), and amplifications were carried out using a C1000 Touch thermocyler (Bio-Rad). Reactions were analyzed using a QX100 droplet reader (Bio-Rad) and quantified using QuantaSoft (v.1.6.6) (Bio-Rad). Results were converted to copy number/g soil extracted by accounting for sample preparation and dilution. For the prepared CaptureSeq libraries, results were converted to copy number/μl by considering dilution factors.

OTU corresponding to the *cpn60* UT plasmids added to the wheat seed wash background were quantified using real-time quantitative PCR (qPCR) primers and amplification conditions as described previously (Dumonceaux et al. 2009). Total bacteria were enumerated using qPCR targeting the 16S ribsosomal RNA-encoding gene as described previously (Lee et al. 1996).

## Results

### 1. CaptureSeq provides microbial community profiles from synthetic microbial ecosystems

#### Zymobiomics reference panel

A simulated microbial community consisting of genomic DNA from 8 bacteria and 2 eukaryotes (with a theoretical composition of 12% each of the 8 bacterial genomes, and 2% each of the 2 eukaryotic genomes) was examined using CaptureSeq, 16S rRNA-encoding gene amplification, and whole metagenome sequencing. To facilitate a direct comparison between 16S amplicon sequencing and CaptureSeq, results were generated using ASV analysis for both methods. Considering only the bacterial genomes that are accessible with 16S amplicon analysis, both methods successfully detected all 8 bacterial OTU (Fig. 1). While the means of the observed proportional composition of the artificial community were identical between the two methods (12.5% across the 8 genomes), CaptureSeq provided a more reproducible profile, with a coefficient of variation of 19% compared to 46.5% for 16S amplification (data not shown). In general, CaptureSeq provided more data from the higher G/C content bacteria compared to 16S amplification, with the highest G/C bacterium (*P. aeruginosa*, 66.2% G/C) found in nearly double the proportional abundance by CaptureSeq compared to 16S amplification. Conversely, *S. aureus* (32.7% G/C) was more accurately represented by 16S amplification compared to CaptureSeq. Overall, however, both methods provided complete and accurate coverage of the bacterial species present in the artificial community.

**Figure 1.**
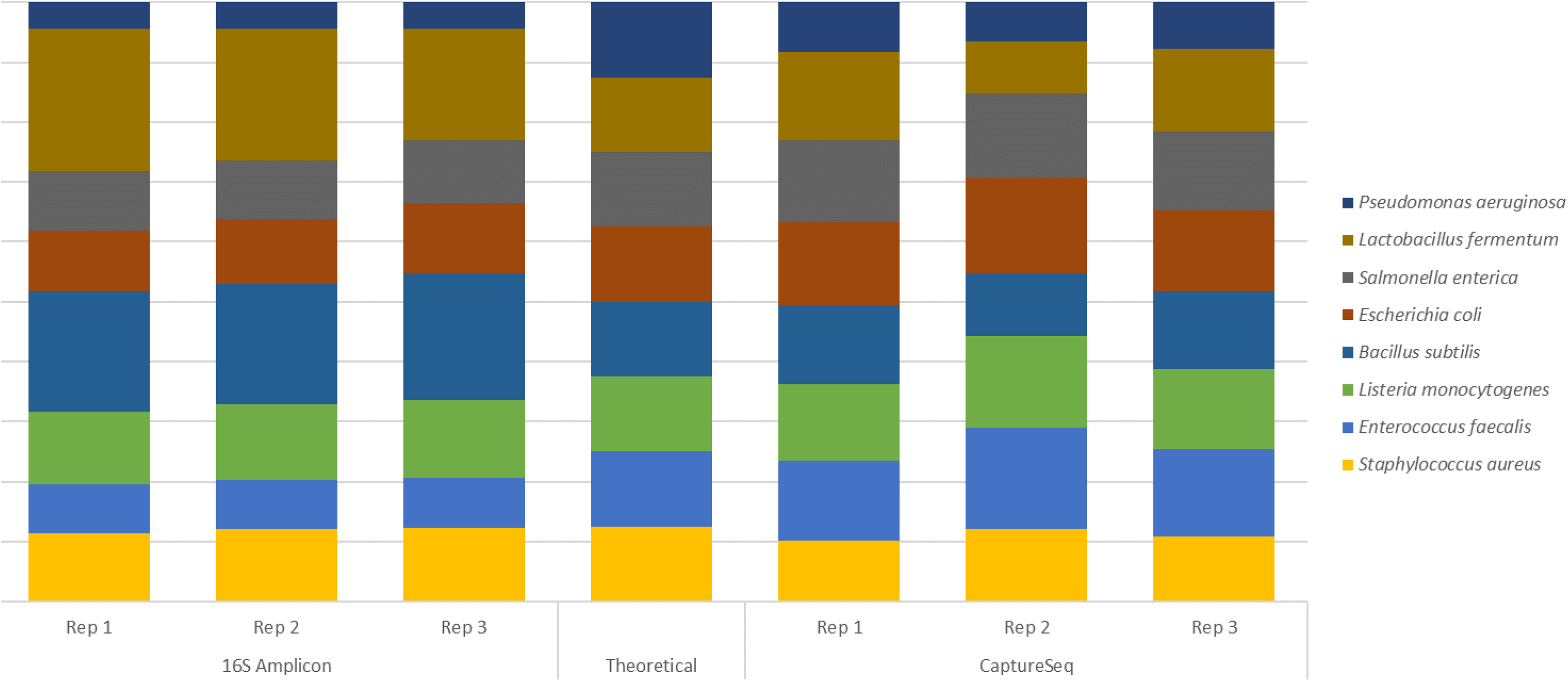
Examination of an artificial microbial community (Zymobiomics Microbial Community Standard) using 16S rRNA-encoding gene amplification and CaptureSeq. The relative abundances of each of the 8 bacterial OTU present in the synthetic community are shown for each of 3 replicates of each method compared to the theoretical composition provided by the manufacturer.

Considering the complete synthetic microbial community, while 16S was unable to identify either of the yeasts present in the mixture, CaptureSeq data examined using ASV analysis identified one of the two yeasts, with ASVs corresponding to *C. neoformans* represented in the CaptureSeq data (Table 1). These ASVs included the beginning of a 64 bp intron found between nucleotides 165 and 166 of the 555 bp *C. neoformans cpn60* universal target (UT) sequence (cpnDB ID b5732). However, no ASVs corresponding to *S. cerevisiae cpn60* were found in the CaptureSeq data.

**Table 1.** Analysis of Zymobiomics microbial community DNA standard using *cpn60*-based CaptureSeq, shotgun sequencing, and taxonomic marker gene amplification.

| organism | CaptureSeq assembly<sup>1</sup> |  |  |  | Shotgun assembly<sup>2</sup> |  |  | sOTU detected<sup>3</sup> |  |
| --- | --- | --- | --- | --- | --- | --- | --- | --- | --- |
|  | *cpn60* UT<br>sequence<br>length | OTU<br>detected | Assembly<br>length<br>(bp) | Sequence<br>identity<sup>4</sup> | *cpn60*<br>OTU<br>detected | Assembly<br>length<br>(bp) | Sequence<br>identity | *cpn60*<br>Capture<br>Seq | 16S<br>amplification |
| **Prokaryotes** |  |  |  |  |  |  |  |  |  |
| *Bacillus subtilis* | 552 | + | 2144 | 100% | + | 873 | 100% | + | + |
| *Escherichia coli* | 555 | + | 1138 | 97% | + | 1208 | 99% | + | + |
| *Enterococcus faecalis* | 552 | + | 1235 | 100% | + | 1133 | 100% | + | + |
| *Lactobacillus fermentum* | 552 | + | 2165 | 100% | + | 870 | 100% | + | + |
| *Listeria monocytogenes* | 552 | + | 1589 | 100% | + | 811 | 99% | + | + |
| *Pseudomonas aeruginosa* | 555 | + | 1183 | 100% | + | 668 | 93% | + | + |
| *Staphylococcus aureus* | 552 | + | 1849 | 100% | + | 487 | 99% | + | + |
| *Salmonella enterica* | 555 | + | 1138 | 97% | + | 1366 | 100% | + | + |
| **Eukaryotes** |  |  |  |  |  |  |  |  |  |
| *Saccharomyces cerevisiae* | 555 | + | 1301 | 100% | + | 384 | 94% | NF<sup>5</sup> | NF |
| *Cryptococcus neoformans* | 619<sup>6</sup> | + | 1435 | 100% | + | 549 | 94% | + | NF |

<sup>1</sup>OTU defined by assembly of CaptureSeq reads using Trinity as described in Methods <sup>2</sup>OTU defined by assembly of shotgun metagenomic reads using Trinity as described in Methods <sup>3</sup>sOTU, sub-OTU as defined by amplicon sequence variant (ASV) analysis using deblur/DADA2 as described in Methods <sup>4</sup>Sequence identity between the assembled OTU and the *cpn60* sequence of the corresponding species. Contains *cpn60* UT and flanking sequences. <sup>5</sup>NF, not found <sup>6</sup>*cpn60* UT of *C. neoformans* contains a 64 bp intron, which was found in the assembled OTU and in the ASV sequence

In addition to ASV analysis, the CaptureSeq data was also used to identify OTU in the artificial community using a reference mapping and *de novo* assembly approach. Using this method, all 8 bacterial *cpn60* sequences and both of the yeast *cpn60* sequences were identified in the CaptureSeq data (Table 1). In every case, the assembled OTU length considerably exceeded the length of the *cpn60* UT that is accessed using the *cpn60* universal PCR primers, and therefore identified the complete UT sequence plus flanking sequences of varying length.

The artificial community was also analyzed using whole metagenome sequencing. This approach identified all of the microorganisms represented in the synthetic microbial community, including all 8 bacteria and both fungi (Table 1). These results were nearly identical to those observed using *cpn60*-based CaptureSeq, except that the assembled fragments tended to be longer and more accurately assembled using CaptureSeq compared to shotgun metagenomic sequencing. (Table 1).

#### Quantification of microbial abundances in CaptureSeq

Using a second synthetic microbial community consisting of 20 *cpn60* UT plasmids spiked into background DNA from a grain wash facilitated a quantitative examination of microbial community profiles generated by CaptureSeq. qPCR quantification of *cpn60* targets from the synthetic community before and after hybridization revealed an enrichment of 3-4 orders of magnitude for *cpn60*-containing DNA fragments compared to 16S rRNA-encoding genes (Table 2). For the 5 exogenous microorganisms that were quantified, the ~10-fold reduction in gene copy number observed between the high, medium, and low spike levels was consistent with the starting composition of the synthetic community samples (Table 2). Furthermore, the number of *cpn60* gene copies for the microorganisms added to the seed wash DNA extract was generally reproducible within each spike level across the 1000-fold difference analyzed. Endogenous microorganisms representing a bacterium (*P. agglomerans*) and a fungus (*Alternaria* sp.) were also detected that corresponded to those previously identified in wheat seed wash samples (Links et al. 2014) (Table 2). Sequencing of the post-hybridization samples revealed a linear correlation between qPCR-determined input gene copies and the number of sequencing reads observed for each of the five targets using the CaptureSeq method, providing Pearson correlation coefficients (r^2^) ranging from 0.995-1.000 (Fig. 2).

**Table 2.** qPCR-determined log_10_ *cpn60* gene copies in wheat seed wash samples spiked with varying amounts of *cpn60* plasmids from non-endogenous microorganisms. Two microorganisms previously shown to be associated with the wheat seed microbiota, along with the total number of 16S gene copies (representing non-*cpn60* DNA) are also shown.

| spike level:<br>microorganism | high |  | medium |  | low |  | unspiked |  |
| --- | --- | --- | --- | --- | --- | --- | --- | --- |
|  | Pre-hyb | Post-hyb | Pre-hyb | Post-hyb | Pre-hyb | Post-hyb | Pre-hyb | Post-hyb |
| *G. vaginalis*<sup>a</sup> | 7.18 | 7.58 | 6.25 | 6.87 | 5.22 | 5.54 | 0.00 | 0.00 |
| *L. crispatus*<sup>a</sup> | 7.20 | 7.49 | 6.26 | 6.81 | 5.28 | 5.45 | 0.00 | 0.00 |
| *L. gasseri*<sup>a</sup> | 7.36 | 8.29 | 6.34 | 7.56 | 5.37 | 6.33 | 0.00 | 0.00 |
| *A. vaginae*<sup>a</sup> | 7.24 | 7.46 | 6.36 | 6.77 | 5.39 | 5.49 | 0.00 | 0.00 |
| *L. iners*<sup>a</sup> | 7.11 | 7.42 | 6.21 | 6.59 | 5.28 | 5.45 | 0.00 | 0.00 |
| *P. agglomerans*<sup>b</sup> | 4.98 | 6.43 | 5.09 | 6.78 | 5.04 | 6.44 | 5.20 | 6.51 |
| *Alternaria sp.*<sup>b</sup> | 5.36 | 4.76 | 5.55 | 5.19 | 5.55 | 4.96 | 5.61 | 5.12 |
| 16S<sup>b</sup> | 8.46 | 5.56 | 8.67 | 5.89 | 8.62 | 5.56 | 8.59 | 5.74 |

<sup>a</sup>exogenous targets (vaginal bacteria *cpn60* plasmids)
<sup>b</sup>endogenous targets (seed wash microorganisms)

**Figure 2.**
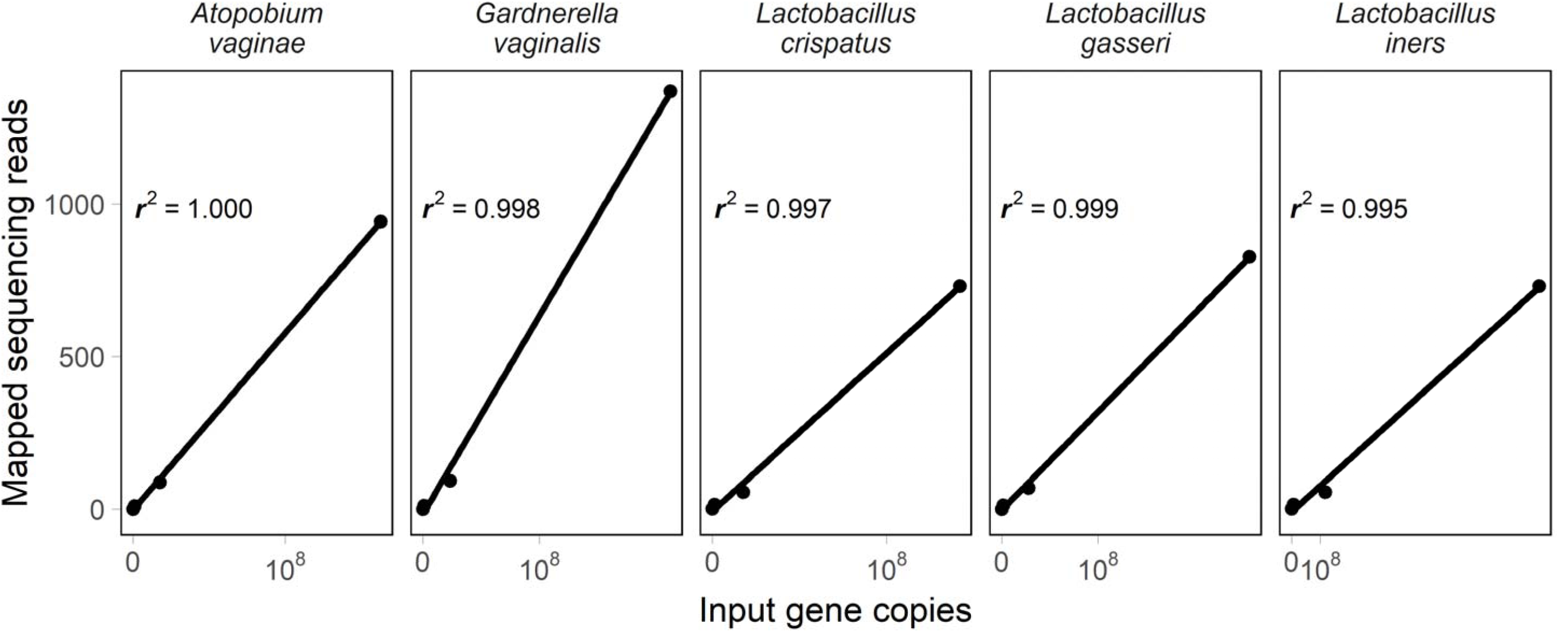
Pearson correlations relating cpn60 gene copies quantified by species-specific quantitative PCR and the number of sequencing reads mapping to each taxonomic cluster by CaptureSeq analysis for 5 bacteria from the synthetic community.

CaptureSeq generated profiles that accurately reflected the relative amounts of DNA spiked into the seed wash background (Table S2). In the CaptureSeq libraries, the number of mapped sequencing reads for each member of synthetic community was within one order of magnitude from the mean for each spike level (Supplemental Fig. S1). *De novo* assembly of the mapped sequencing reads for each microorganism from the *cpn60* plasmid synthetic panel generated OTU that were >99% identical to the known *cpn60* sequences (not shown). Based on the results observed using these synthetic microbial communities, analysis of natural microbial communities used the reference mapping and *de novo* assembly approach. To facilitate comparison, microbial communities from natural microbial ecosystems used CaptureSeq and whole metagenome sequencing.

### 2. CaptureSeq provides microbial community profiles in natural ecosystems

Microbial profiles were generated by CaptureSeq and whole metagenome sequencing using samples from natural environmental ecosystems including soil, manure storage tanks, and a non-aerated terrestrial pond. These samples were chosen to reflect high complexity and principally Bacteria (soil), samples enriched in Archaea (manure storage tank), and samples with higher numbers of Eukarya (freshwater pond). The CaptureSeq profiles of these communities provided a taxonomic overview of Bacteria, Archaea and Eukarya simultaneously, and contained sequencing reads from 9,361 (soil), 9,306 (manure), and 6,568 (pond) distinct taxonomic clusters (Supplemental Dataset S3). Additionally, the CaptureSeq profiles facilitated inter-Domain comparisons of read abundances among taxonomic groups, since the abundances could be expressed in relation to the total pan-Domain community as opposed to reflecting only the proportions within a single Domain (Fig. 3).

**Figure 3.**
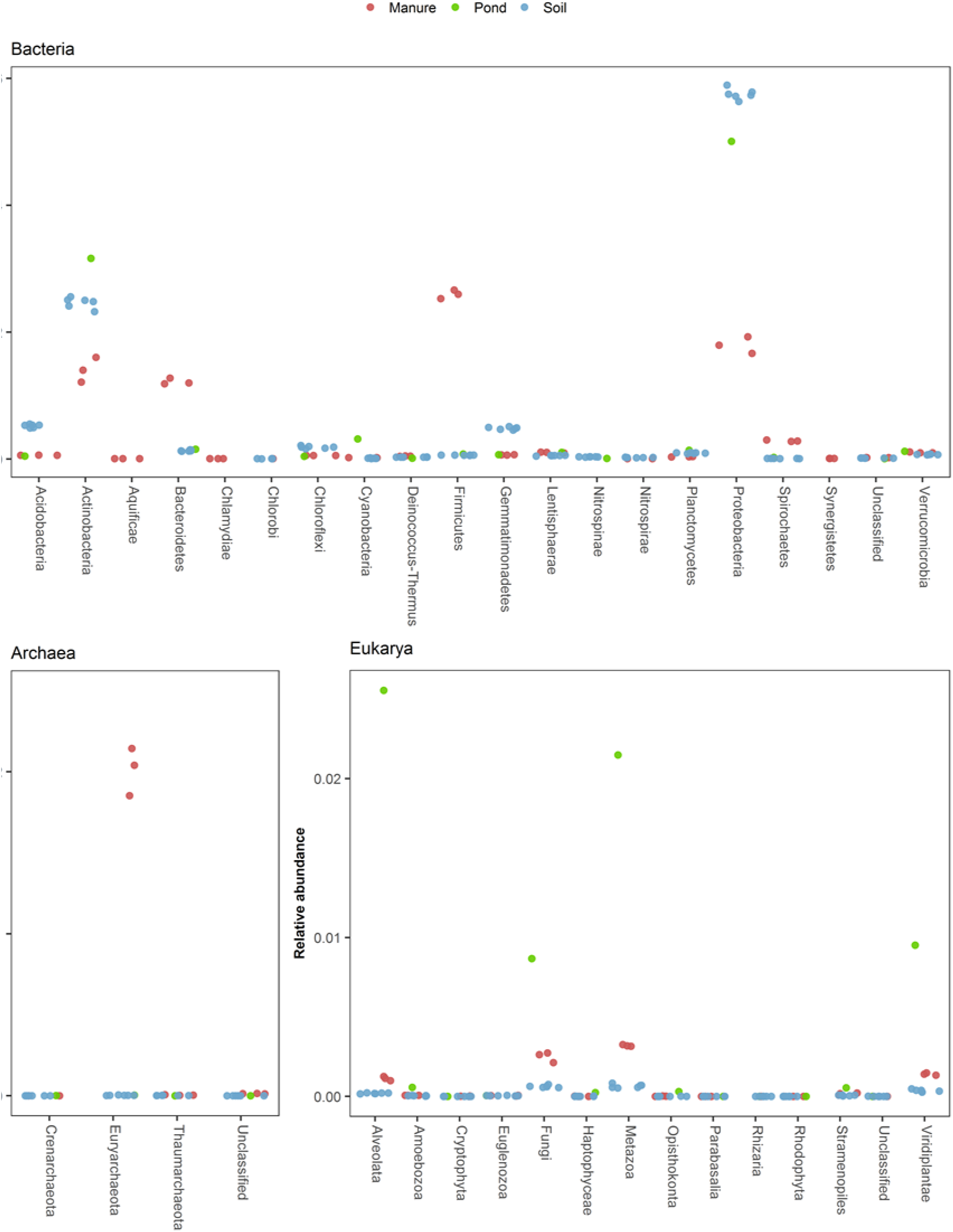
CaptureSeq results obtained by reference mapping on an ecologically diverse range of samples including soil (n=6); manure storage tanks (n=3); and a freshwater pond (n=1). The relative abundances of individual phyla are expressed as a proportion of the entire pan-Domain microbial communities.

The soil sample microbiomes, as expected, were composed primarily of Bacteria, with Proteobacteria and Actinobacteria comprising 60% and 25% of the pan-Domain community respectively. Consistent with observations in the synthetic communities, high G/C Actinobacteria were well represented in the microbial community profiles generated by CaptureSeq for the soil samples, including several members of the genera *Nocardiodes, Marmoricola*, and *Pseudonocardia* with G/C contents ranging from 64-71% (not shown). Members of the phyla Acidobacteria and Gemmatinomonadetes represented an additional 5% each of the soil microbiome. Total archaeal reads only accounted for 0.03-0.08% of the soil microbial community, however there were still 165 archaeal taxonomic clusters identified in the soil. Eukarya represented just 0.18-0.21% of the soil microbiome, with Fungi and Metazoa the most abundant taxonomic groups. While the bovine manure storage tank-derived samples also contained a diverse array of Bacteria, they only represented 77-80% of the microbiome, compared to >99% for the soil samples. CaptureSeq libraries from the manure storage tank samples contained 19-22% archaeal reads, of which the vast majority were methanogens from the Phylum Euryarchaeota. The terrestrial pond contained a much greater proportion and diversity of Eukaryotes, representing 6.7% of the sequencing reads and 361 taxonomic clusters (Supplemental Dataset S3).

### 3. *CaptureSeq data facilitates the assembly of target OTU from taxonomic clu*sters

Analysis of the synthetic microbial communities demonstrated that the *cpn60* molecular barcode could be reconstituted from CaptureSeq data using *de novo* assembly of OTU. To determine if the assembly of OTU representing individual organisms was reliable in complex natural microbial communities using CaptureSeq, we selected target taxonomic clusters identified through reference mapping for assembly. *De novo* assembly of eukaryotic sequencing reads from the terrestrial pond sample generated 11 OTU most closely related to members of the Phylum Chlorophyta (green algae). Additionally, the assembly of OTU most similar to *Aenopholes* sp. (mosquitoes), and three members of the Phylum Alveolata (protists), suggested that CaptureSeq was able to retrieve *cpn60* DNA from a diverse array of Eukarya. Compared to reference sequences in cpnDB, these *de novo* assembled OTU had nucleotide identities ranging from 59-84%, suggesting that the current probe array design and hybridization conditions were sufficiently permissive to allow the capture of novel *cpn60* sequences (true unknowns).

To examine the suitability of *de novo* assembly for providing taxonomic markers suitable for tracking the abundances of particular OTU across samples, we selected target microorganisms for quantification in antibiotic-treated soil samples using OTU-specific qPCR. For Bacteria, we quantified *Microbacterium lacus* strain C448, which was previously cultured from these soil samples and shown to degrade and metabolize the sulfonamide antibiotic added to the field plots (Topp et al. 2013). While the presence of this target in the soil samples was confirmed using culture methods, it was under-represented in the shotgun metagenomic libraries when compared to the CaptureSeq profiles. Only the CaptureSeq data provided a sufficient number of target sequencing reads for *de novo* assembly, generating a 1,066 bp OTU that was >99% identical to the *cpn60* sequence obtained from the genome of this organism (Martin-Laurent et al. 2014). Quantification of *M. lacus* C448 showed that the bacterium was present at a low level in all soil samples of between 10^3^ and 10^4^ gene copies per gram of soil, and that the levels were significantly higher in the 10 ppm antibiotic-treated soil samples compared to untreated soils (Table 3).

**Table 3.** Abundances of selected OTU from each Domain in antibiotic treated soil samples, as determined by quantitative PCR.

| OTU | Domain | cpnDB nearest neighbor | OTU Length (bp) | Sequence identity (%)<sup>1</sup> | Treatment (mg kg<sup>-1</sup>) | soil extract (copies/g soil) | Post-hybridization sample (copies/μl) |
| --- | --- | --- | --- | --- | --- | --- | --- |
| XP002901426<br>DN2_c0_g1_i1 | Eukarya<br>(type I)<sup>2</sup> | *Phytophthora infestans* | 539 | 100 | 0<br>10 | ND<sup>3</sup><br>ND | 1242<br>3942 |
| WP036300323<br>DN4_c3_g1_i2 | Bacteria<br>(type I) | *Microbacterium lacus C448* | 1,066 | 99 | 0<br>10 | 6750<br>38571<sup>4</sup> | 1417<br>8170<sup>4</sup> |
| KUL05486<br>DN0_c0_g1_i1 | Archaea<br>(type II) | *Methanoculleus marisnigri* | 1,029 | 92 | 0<br>10 | 495<br>527 | ND<br>3360 |

<sup>1</sup>Percent identity to reference sequence in cpnDB.
<sup>2</sup>Type I refers to the ~60 kDa mitochondrial and chloroplast proteins found in Bacteria, Eukarya, and certain Archaea. Type II refers to TCP1, the cytoplasmic orthologue of the group I chaperonins found in Archaea.
<sup>3</sup>ND, not detected.
<sup>4</sup>Statistically significant difference ( $p < 0.01$ ) between 0 mg kg<sup>-1</sup> and 10 mg kg<sup>-1</sup> groups, using a Mann-Whitney rank sum test

A *cpn60* OTU corresponding to *Acinetobacter baumanii/A. calcoaceticus* was also assembled from the CaptureSeq data and quantified using ddPCR. This OTU was determined by CaptureSeq to be maximally abundant in plot 4, which was 1 of 3 replicates that had been treated with 10 ppm of antibiotics. Consistent with the CaptureSeq data, ddPCR revealed that the OTU corresponding to *A. baumanii* had by far the highest abundance in plot 4 (approximately 10^6^ genomes/g soil) and was present at levels near or below 10^3^ genomes/g soil in other soil samples in which reads corresponding to this OTU were not detected (Fig. 4). Plots 11 (10 ppm) and 3 (0.1 ppm) also had *A. baumanii* OTU in the CaptureSeq datasets and this OTU was present at slightly higher levels in these plots (Fig. 4).

**Figure 4:**
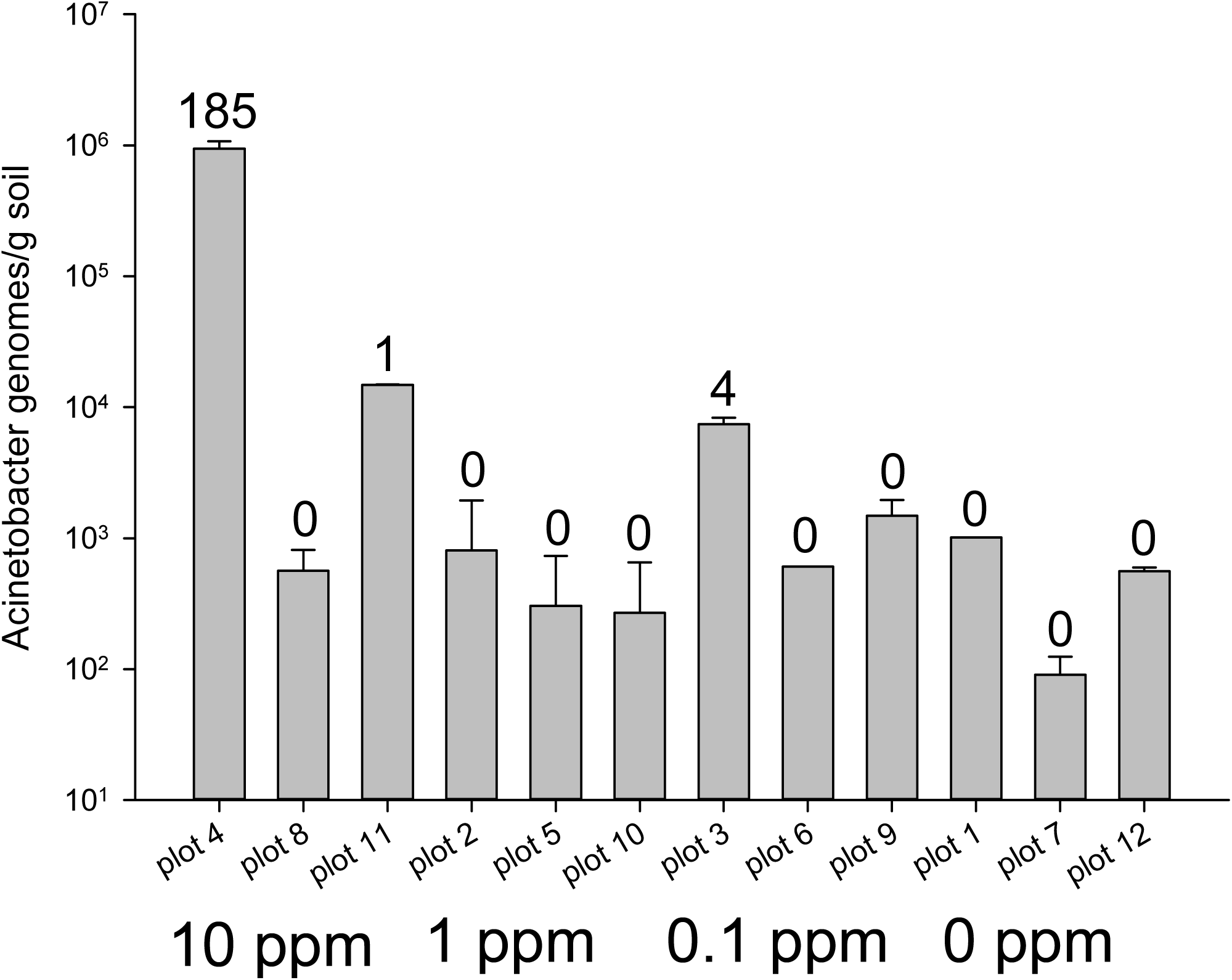
Quantification of assembled OTU DN9_c0_g1_i1, corresponding to *Acinetobacter baumanii/A. calcoaceticus*, in soil samples treated with antibiotics. The number of sequencing reads mapping to this taxonomic cluster in each soil sample is indicated above each bar.

Assembled OTU targets from the Domains Eukarya (type I-*Phythophthora infestans*) and Archaea (type *II-Methanoculleus* sp.). were also selected for quantification using ddPCR. The archaeal OTU was quantified at levels between 495 and 527 gene copies per gram of soil. The OTU corresponding to *P. infestans* was present at levels below the limit of detection of ddPCR for these samples, yet was detectable by CaptureSeq (Table 3). These results suggest that the CaptureSeq method could nearly completely sample complex microbial communities with a limit of detection beyond the dynamic range of even very sensitive quantification methods like ddPCR.

### 4. CaptureSeq requires significantly less sequencing input compared to whole metagenome sequencing to achieve adequate community coverage

The number of sequencing reads corresponding to *cpn60* genes represented 0.07% of the total reads from the whole metagenome library compared to an average of 16.7% (± 0.8%) for CaptureSeq (Table S3). Examining the Good’s coverage estimator (Chao et al. 2015) as a function of sequencing depth revealed that CaptureSeq provided almost complete community coverage in soil while whole metagenome sequencing required orders of magnitude greater sequencing effort to achieve a high level of community coverage (Fig. 5).

**Figure 5:**
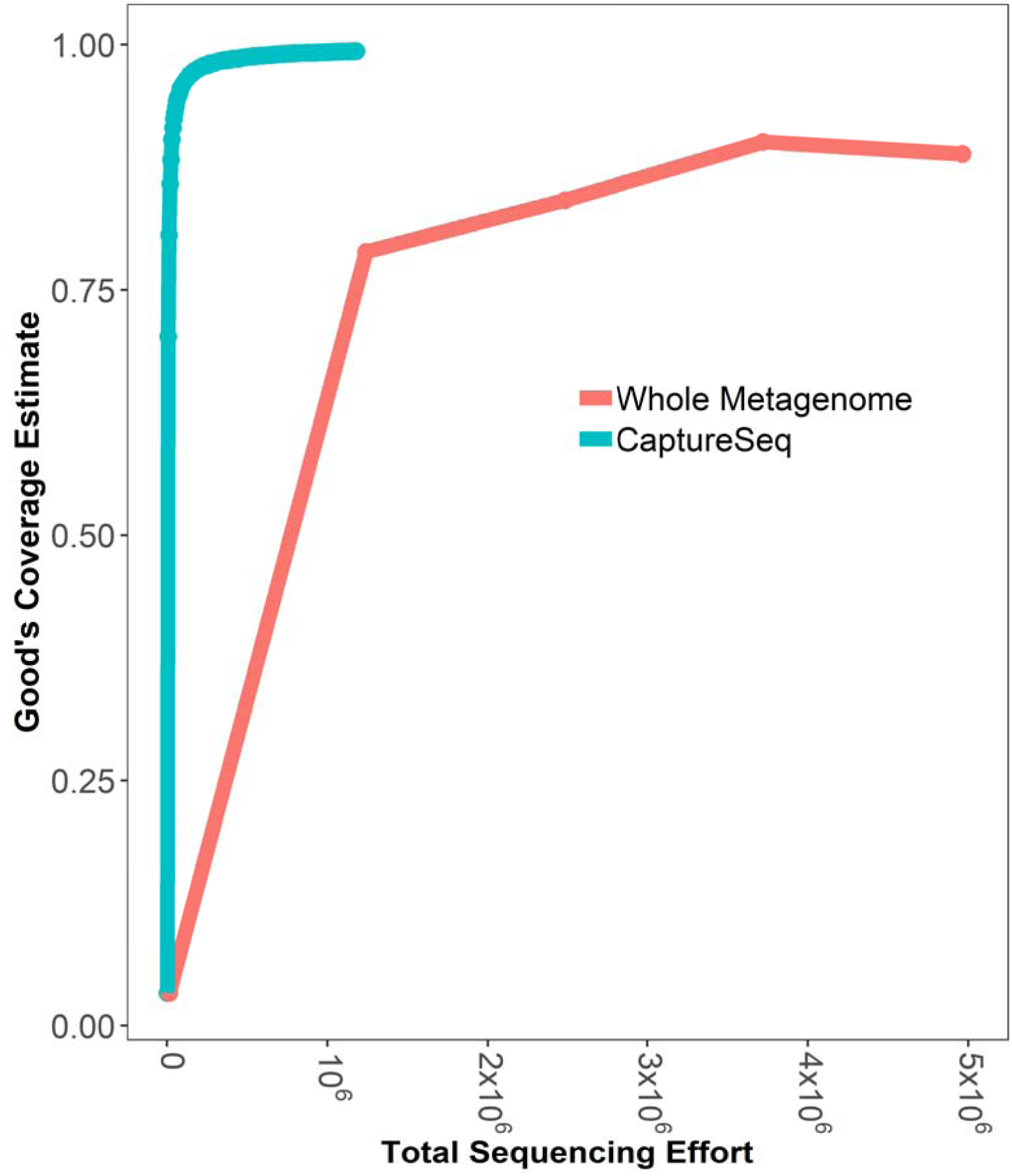
Good’s coverage estimate reflecting the average total sequencing effort for six soil samples each profiled using CaptureSeq (blue), or whole metagenome sequencing(red) approaches.

### 5. Microbial ecosystem diversity of soil samples as determined by CaptureSeq and whole metagenome sequencing

Examination of OTU abundance patterns using hierarchical clustering comparing both methods (inter-method) revealed that the samples displayed patterns of microbial abundances that clustered strongly by analysis method (Fig. 6). Similarly, intra-method beta diversity assessed using the Bray-Curtis dissimilarity metric revealed that while the communities clustered strongly by method (Fig. 7A), CaptureSeq uniquely suggested a possible treatment effect of the 10 ppm antibiotic in the long-term treated soil samples (Fig. 7 B,C).

**Figure 6.**
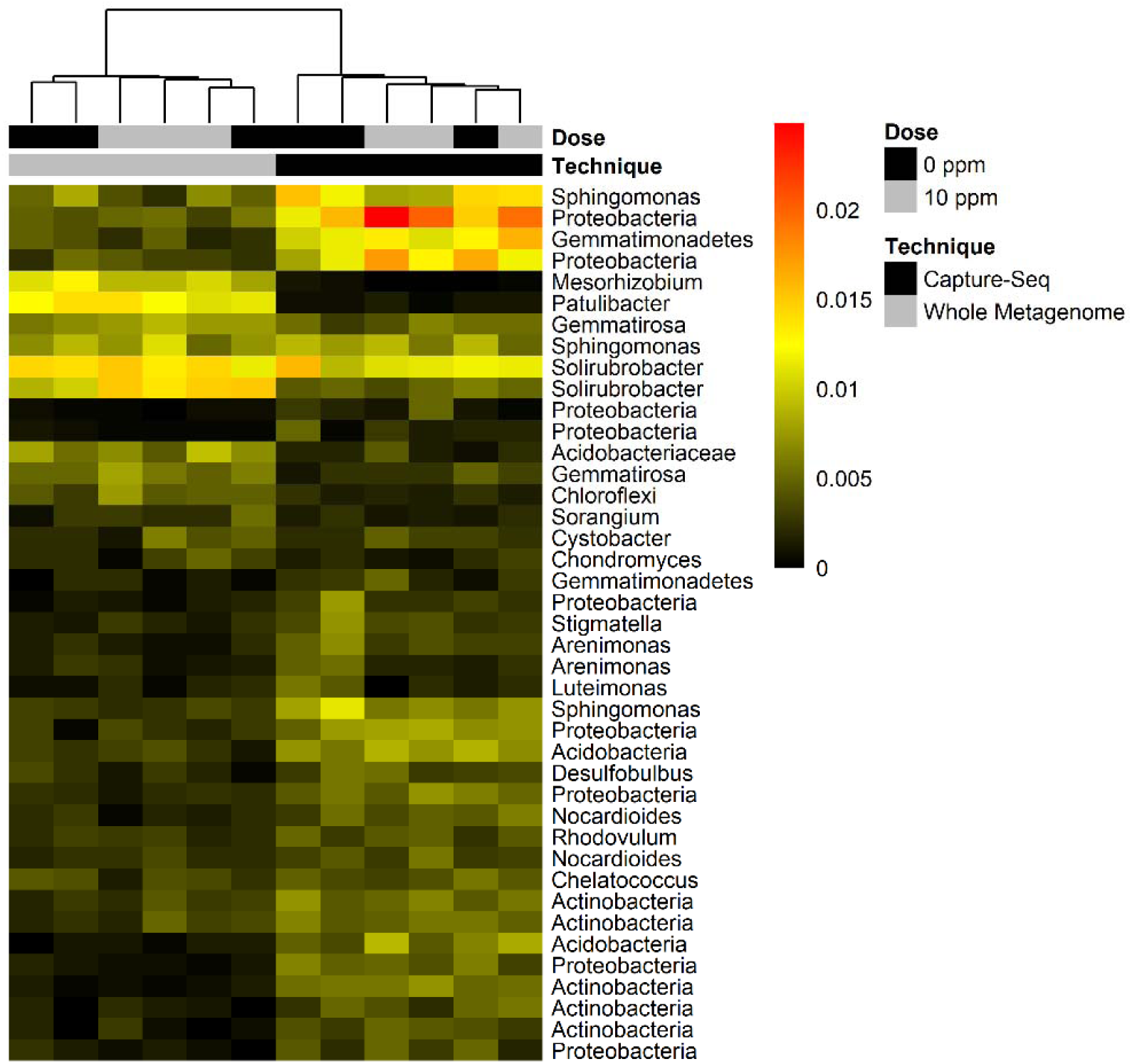
Proportional abundances of taxonomic clusters obtained by reference mapping for type I chaperonins in soil samples profiled using CaptureSeq or shotgun metagenomic approaches. Samples were clustered based on Bray-Curtis distance, and reference clusters composing a minimum of 0.5% of the mapped sequencing reads in any one sample are shown. Samples are coded according to antibiotic treatment: black (10 ppm) or gray (0 ppm).

**Figure 7.**
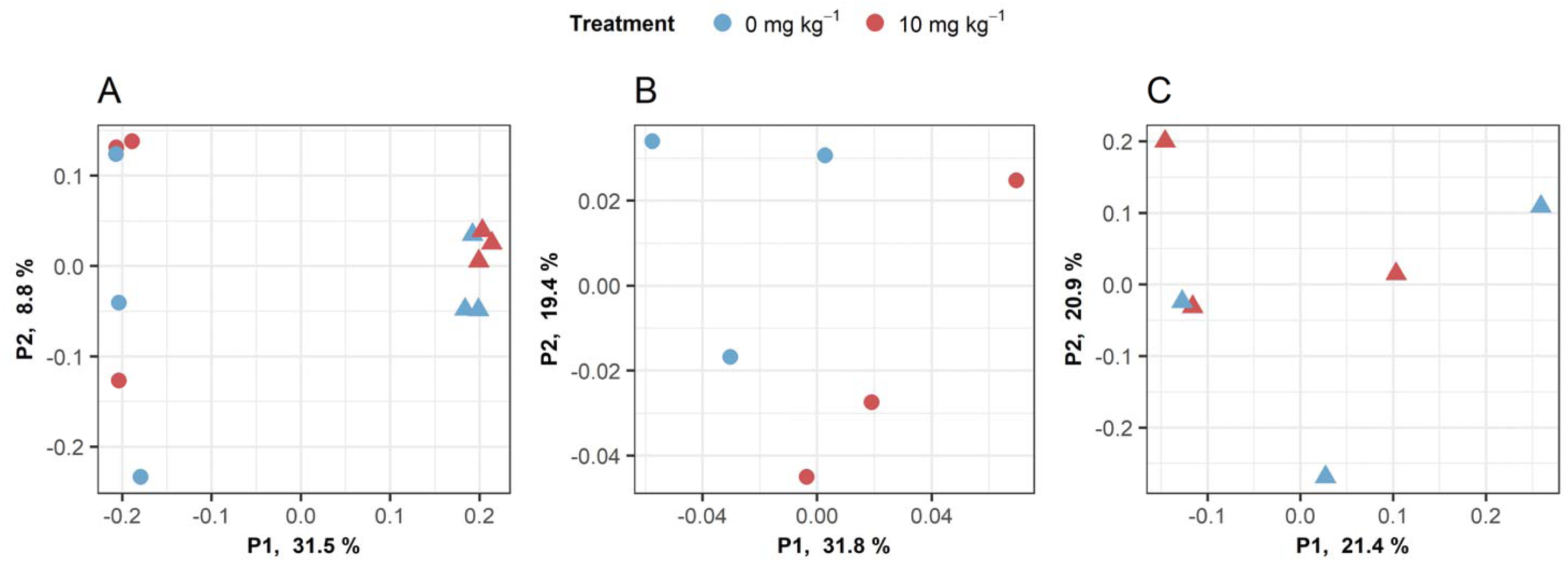
Principal coordinate analysis of Bray-Curtis dissimilarity between soil samples profiled using CaptureSeq (circle) or whole metagenome sequencing (triangle) approaches. OTU frequencies were determined by reference mapping as described in the text. **A.** Both approaches considered together. **B.** CaptureSeq. **C.** Whole metagenome sequencing.

Comparing alpha diversity metrics of the soil communities between the profiling techniques, richness (Chao1) (p = 0.004), evenness (Simpson 1-D) (p = 0.004) and diversity (Shannon H’) (p = 0.004) were all higher when profiled using whole metagenome sequencing compared to CaptureSeq (Fig. 8). Additionally, the Simpson (p = 0.034) and Chao1 (p = 0.012) diversity metrics of the CaptureSeq method showed lower variance among the biological replicates of each treatment, even when libraries were down-sampled to very low levels (Fig. 8).

**Figure 8:**
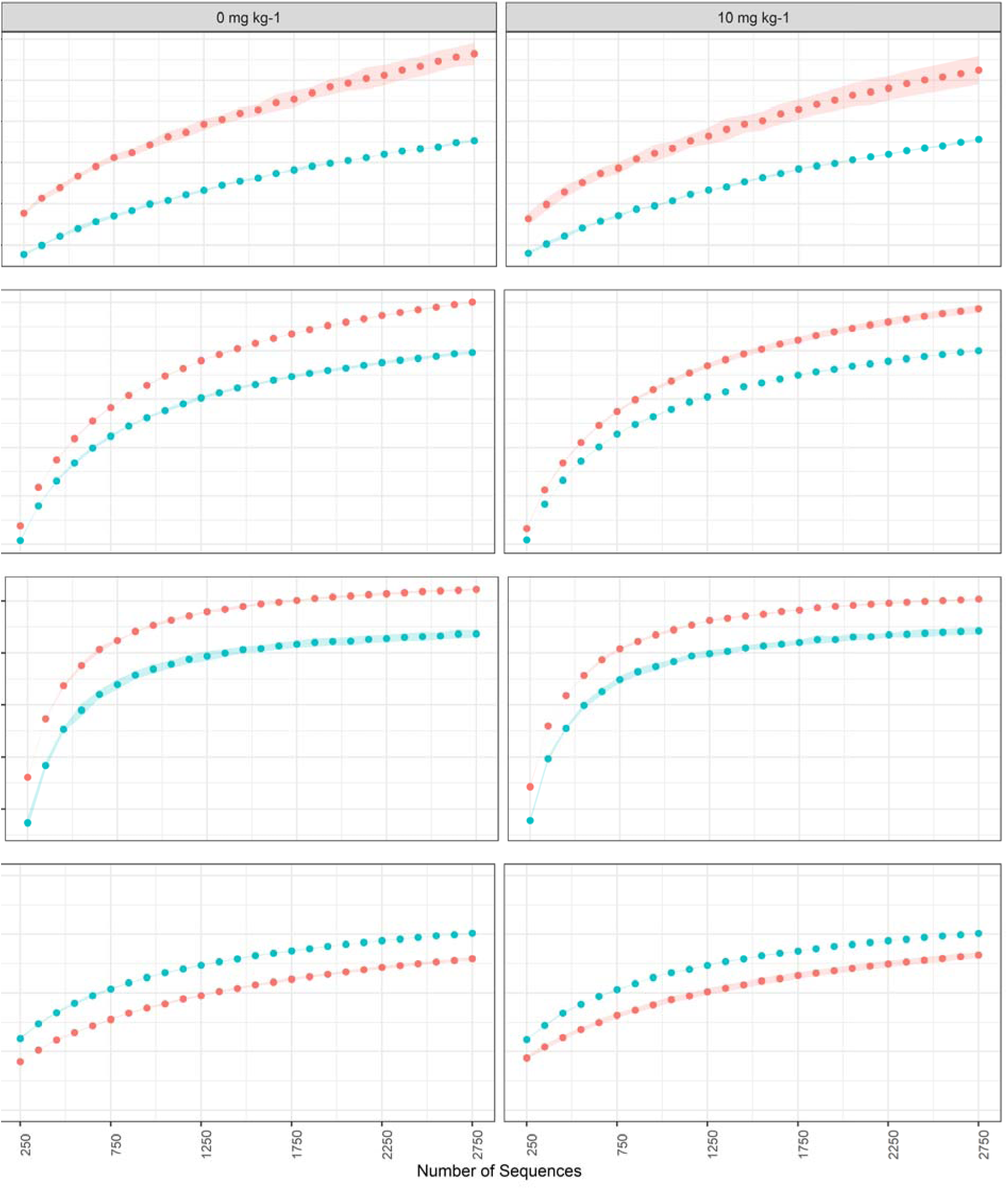
Alpha diversity metrics for soil samples profiled using CaptureSeq (blue) or whole metagenome sequencing (red) approaches. Metrics were calculated using libraries that were downsampled from 250-2750 reads and were averaged across 100 bootstrapped datasets. The shaded area corresponds to the standard deviation of the three replicate soil plots for each antibiotic treatment condition.

## Discussion

Experimental approaches to determining the taxonomic composition of microbial ecosystems have typically employed PCR amplification of Domain-restricted taxonomic markers, typically various regions of the 16S rRNA-encoding gene for Bacteria (Caporaso et al. 2018) and the ITS locus for Fungi (Schoch et al. 2012). In addition, PCR amplification of the *cpn60* UT can generate microbial community profiles that include bacteria and eukaryotes simultaneously (Links et al. 2014). While these methods can provide a rapid and cost-effective means of determining the taxonomic composition of a microbial community, they also have well recognized limitations associated with low taxonomic resolution and PCR amplification biases (Edgar 2017; Knight et al. 2018). These drawbacks can be partially overcome with whole metagenome shotgun sequencing (Ranjan et al. 2016), but this approach is not feasible in many experimental models. These considerations have led to the development of several alternative means of profiling microbial communities, such as 16S rRNA-encoding gene-based hybridization capture (Gasc & Peyret 2018), transfer RNA sequencing (Schwartz et al. 2018), and real-time single molecule sequencing of full-length 16S rRNA-encoding genes (Singer et al. 2016). Each of these methods presents unique advantages aimed at overcoming some of the limitations of PCR-based profiling, but each method also comes with its own limitations that must be considered when deciding on an appropriate method for a particular experimental system.

Here we have described an experimental approach for determining the taxonomic composition of microbial communities that exploits the features of solution-based hybridization of *cpn60* sequences. This approach utilizes the advantages of *cpn60*-based microbial community profiling, including its high taxonomic resolution and ability to profile multiple Domains simultaneously (Links et al. 2014; McKenney et al. 2014; Peterson et al. 2016; Town et al. 2014). While *cpn60* has proven to be a powerful and useful taxonomic marker for microbial community analysis, PCR amplification of *cpn60* targets in microbial communities presents analogous biases in community representation as seen in 16S and ITS-based amplification strategies (Hill et al. 2005).

This work provides experimental and computational methods for analyzing microbial community composition using *cpn60*-based hybridization and demonstrates that CaptureSeq provides quantitatively reliable data that can detect OTU from multiple Domains within the same dataset. Nevertheless, improvements to both the experimental and computational parts of the workflow can be considered. On the experimental side, the composition of the hybridization array could be optimized and expanded based on the accumulation of novel *cpn60* sequences from currently unrepresented taxa. This would help to minimize biases associated with the lack of representation of taxa with no probes on the capture array of sufficient nucleotide sequence similarity. The hybridization conditions could also potentially be optimized to maximize the recovery of *cpn60* sequences in the dataset. The current workflow resulted in an approximately 200-fold enrichment of the soil samples for the taxonomic marker of interest, from under 0.1% of reads in the shotgun metagenomic sequencing to over 15% of reads in the CaptureSeq datasets. While this level of enrichment enabled deep sampling of the soil microbial communities (similar to that attained using PCR-based enrichment-not shown), it does not approach the levels of enrichment that were observed using 16S-based hybridization (Gasc & Peyret 2018). More recent CaptureSeq experiments employing newer blocking oligonucleotides provided by the manufacturer has resulted in improvements to the recovery of *cpn60* gene fragments – up to 70% of reads in recent studies (data not shown).

Computationally, a key challenge to be overcome in the analysis of CaptureSeq data, like PCR amplification-based methods, is the most appropriate means of defining an OTU. We have demonstrated here that the assembly of *cpn60* gene fragments exceeding even the length of the 549-567 bp *cpn60* UT (Hill & Vancuren 2019) is possible using CaptureSeq data, but assembly algorithms can be computationally challenging on large datasets, and the formation of artefactual OTU can be difficult to avoid. Targeted assembly of specific OTU from reference bins obtained by mapping offers a means to overcome these difficulties and can provide taxonomic markers suitable for strain tracking across experimental systems and for strain culture. Moreover, other assembly methods are available that do not require prior reference binning; for example, we have successfully assembled *cpn60* OTU from complex CaptureSeq data without reference mapping using TransAbyss (Robertson et al. 2010). Nevertheless, alternative means of defining OTU have been described for 16S PCR amplicon data that exploit unique sequence variants and are recommended for nearly all applications (Knight et al. 2018). CaptureSeq can be used to define analogous *cpn60*-based sub-OTU (sOTU) using DADA2 (Callahan et al. 2016); however, the read length of Illumina data precludes the use of the entire ~550 bp *cpn60* UT sequence for OTU definition. Recent work has determined that the first 150 bp of the *cpn60* UT is suitable for sOTU definition (Vancuren et al. 2020). However, the ASV method appeared to have a lower sensitivity compared to assembly, as it failed to detect one of the eukaryotic OTU from the Zymobiomics reference panel. Further refinement of sOTU definition is a requirement for future CaptureSeq studies. We suggest that the sOTU approach may be suitable for experimental questions aimed at determining and comparing total microbial community structure across samples, while the assembly approach may be more suitable for experimental questions aimed at identifying particular microorganisms that may be associated with some desired function – this is because the assembly approach provides more taxonomic information for microorganism identification (by culture) and tracking across experimental systems due to the length of the assembled *cpn60* OTU.

While the effects of long-term antibiotic treatments on soil microbial communities was not a focus of this study, we observed broad differences in the taxonomic composition and diversity of microbial communities in soil samples treated with antibiotics using CaptureSeq compared to untreated soil, with antibiotic-treated communities clustering distinctly from the untreated soil samples. The strong correlations we observed between OTU abundances and qPCR-determined abundances in CaptureSeq data in the synthetic panel, along with the agreement between CaptureSeq read frequencies and ddPCR-determined abundances for the bacteria analyzed in the natural soil ecosystem (*Microbacterium* and *Acinetobacter*), suggest that the CaptureSeq data accurately represented the composition of these microbial communities. The fact that known antibiotic-degrading bacteria (*M. lacus*) were detected in higher abundances in the antibiotic treated soil is consistent with the selection of such bacteria in the presence of antibiotics. In addition, the increased abundance of *A. baumanii* in antibiotic treated soils, which was noted by CaptureSeq and by ddPCR, is consistent with the known ability of this organism to acquire antibiotic resistance genes through horizontal gene transfer (Cooper et al. 2017).

While the experimental and analytical methods for CaptureSeq could be improved, the method offers several distinct advantages compared to PCR amplification and shotgun metagenomics for analyzing microbial communities. For example, CaptureSeq provided a balanced view of the relative abundances of microorganisms within the community. PCR-associated representational bias, which presents a skewed representation of microbial taxon abundance (Props et al. 2016), is a well-known phenomenon (Green et al. 2015; Johnson et al. 2015; Lee et al. 2012). CaptureSeq also resulted in an improvement of the representation of high G/C content microorganisms compared to amplification. Difficulty in amplification of high G/C content targets is a phenomenon that has been previously observed using both 16S and *cpn60* taxonomic markers from mixed communities (Hill et al. 2006; Pinto & Raskin 2012). *De novo* assembly of taxonomic clusters from the CaptureSeq datasets into OTU for which probes were not explicitly designed, such as *M. lacus* strain C448, also suggests that off-target *cpn60* sequence capture can expand the breadth of OTU observed in the dataset beyond the sequences represented in the probe array and can include sequences that have not been previously observed. Off-target hybridization resulting in the identification of novel taxa was also observed using 16S rRNA gene-based hybridization (Gasc & Peyret 2018).

Both CaptureSeq and whole metagenome sequencing provided the means to identify OTU from all Domains simultaneously, facilitating the characterization of inter-Domain relationships among microorganisms. The ability to calculate the abundances of organisms as a proportion of the entire pan-Domain community facilitates the identification of inter-Domain relationships and syntrophies. This is of particular importance in many settings (e.g. manure or gut health) in identifying the syntrophic relationships between volatile fatty acid producing Bacteria and methanogenic Archaea (Demirel & Scherer 2008). In soil, the complex relationship between saprophytic Fungi and Bacteria is critical to examining the role of the microbiome in nutrient cycling (de Menezes et al. 2017). This advantage is not offered using amplification of universal targets, although PCR-based enrichment does provide the benefit of very deep coverage of complex microbial communities. Whole metagenome sequencing does not provide the community coverage of the CaptureSeq method at a similar sequencing effort, suggesting that complex microbiomes will likely require additional phylogenetic data to make any informed examination of microbial diversity metrics. Whole metagenome sequencing can reasonably be considered to be a less biased means of determining the taxonomic composition of an environmental sample, and may be a suitable choice when sufficient sequencing resources are available. However the abiding popularity of amplicon-based profiling is at least partially a result of the high degree of enrichment of taxonomically informative sequence reads that it generates. CaptureSeq provides an alternative that avoids the amplification biases associated with PCR while retaining the sequencing efficiency of amplicon-based profiling.

Molecular microbial community profiling is one of the foundational steps in exploring microbiome structure-function relationships in an experimental system (Carballa et al. 2015; Gopal & Gupta 2016; Muegge et al. 2011). To generate and evaluate scientific hypotheses it is critical to generate a microbiome profile that reflects the natural state as closely as possible with sufficient sensitivity to evaluate both abundant and rare microorganisms. The *cpn60*-based method described herein permits taxonomically broad and deep microbial community profiling of complex microbiomes. Thus CaptureSeq has the potential to impact life sciences research wherever microbes are thought to be important, including human health and nutrition (Kau et al. 2011), agriculture (Busby et al. 2017), biotechnology (Koch et al. 2014), and environmental sciences (Fierer 2017). In each of these areas, researchers can choose from an increasing array of tools to address the particular experimental question at hand. While all microbial community profiling techniques have inherent limitations and biases, CaptureSeq is a suitable alternative that provides quantitative, cross Domain data for the analysis of complex microbial ecosystems.

## Conclusions

In this work, we have demonstrated the utility of *cpn60*-based hybridization to enrich environmental DNA samples for taxonomically informative DNA sequences. Using synthetic microbial ecosystems, CaptureSeq was shown to provide robust results regarding the presence and abundance of prokaryotes and eukaryotes simultaneously, and that the read abundances generated are quantitatively reliable. CaptureSeq was also shown to provide pan-Domain microbial community profiles in complex natural ecosystems and generate read abundances that are consistent with observations made using microorganism-targeted molecular diagnostic assays. While both the experimental and computational methods could be improved, this work represents an important first step in demonstrating the utility of this approach for providing microbial community profiles that reflect the natural state more closely than PCR-based methods. CaptureSeq could easily be applied to a wide range of microbial ecosystems, providing robust data on microorganism presence and abundance that can inform many experimental questions regarding the role of the microbiota in human, environmental, and agricultural ecosystems.

## Supporting information

Supplemental file pseudocode

## Acknowledgements

We thank Christine Hammond for technical improvements to the CaptureSeq method and for valuable commentary on this manuscript.

## References

Adékambi T, Drancourt M, and Raoult D. 2009. The *rpoB* gene as a tool for clinical microbiologists. Trends Microbiol 17:37–45.

Barret M, Briand M, Bonneau S, Préveaux A, Valière S, Bouchez O, Hunault G, Simoneau P, and Jacques M-A. 2015. Emergence shapes the structure of the seed-microbiota. Appl Environ Microbiol 81:1257–1266.

Barret M, Gagnon N, Kalmokoff ML, Topp E, Verastegui Y, Brooks SPJ, Matias F, Neufeld JD, and Talbot G. 2013. Identification of *Methanoculleus* spp. as active methanogens during anoxic incubations of swine manure storage tank samples. Appl Environ Microbiol 79:424–433.

Busby PE, Soman C, Wagner MR, Friesen ML, Kremer J, Bennett A, Morsy M, Eisen JA, Leach JE, and Dangl JL. 2017. Research priorities for harnessing plant microbiomes in sustainable agriculture. PLOS Biology 15:e2001793. 10.1371/journal.pbio.2001793

Callahan BJ, McMurdie PJ, Rosen MJ, Han AW, Johnson AJA, and Holmes SP. 2016. DADA2: High-resolution sample inference from Illumina amplicon data. Nature Meth 13:581. 10.1038/nmeth.3869 https://www.nature.com/articles/nmeth.3869#supplementary-information

Caporaso JG, Ackermann G, Apprill A, Bauer M, Berg-Lyons D, Betley J, Fierer N, Fraser L, Fuhrman JA, Gilbert JA, Gormley N, Humphrey G, Huntley J, Jansson JK, Knight R, Lauber CL, Lozupone CA, McNally S, Needham DM, Owens SM, Parada AE, Parsons R, Smith G, Thompson LR, Thompson L, Turnbaugh PJ, Walters WA, and Weber L. 2018. EMP 16S Illumina Amplicon Protocol. protcolsio. http://dx.doi.org/10.17504/protocols.io.nuudeww

Caporaso JG, Kuczynski J, Stombaugh J, Bittinger K, Bushman FD, and Costello EK. 2010. QIIME allows analysis of high-throughput community sequencing data. Nat Methods 7. 10.1038/nmeth.f.303

Carballa M, Regueiro L, and Lema JM. 2015. Microbial management of anaerobic digestion: exploiting the microbiome-functionality nexus. Curr Opin Biotechnol 33:103–111. http://dx.doi.org/10.1016/j.copbio.2015.01.008

Chaban B, and Hill JE. 2012. A ‘universal’ type II chaperonin PCR detection system for the investigation of Archaea in complex microbial communities. ISME J 6:430–439.

Chao A, Hsieh TC, Chazdon RL, Colwell RK, and Gotelli NJ. 2015. Unveiling the species-rank abundance distribution by generalizing the Good-Turing sample coverage theory. Ecology 96:1189–1201. 10.1890/14-0550.1

Cleary DW, Bishop AH, Zhang L, Topp E, Wellington EM, and Gaze WH. 2016. Long-term antibiotic exposure in soil is associated with changes in microbial community structure and prevalence of class 1 integrons. FEMS Microbiol Ecol 92. 10.1093/femsec/fiw159

Cooper RM, Tsimring L, and Hasty J. 2017. Inter-species population dynamics enhance microbial horizontal gene transfer and spread of antibiotic resistance. eLife 6:e25950. 10.7554/eLife.25950

de Menezes AB, Richardson AE, and Thrall PH. 2017. Linking fungal–bacterial co-occurrences to soil ecosystem function. Curr Opin Microbiol 37:135–141. http://dx.doi.org/10.1016/j.mib.2017.06.006

Demirel B, and Scherer P. 2008. The roles of acetotrophic and hydrogenotrophic methanogens during anaerobic conversion of biomass to methane: a review. Rev Environ Sci Biotechnol 7:173–190.

Dumonceaux TJ, Hill JE, Hemmingsen SM, and Van Kessel AG. 2006. Characterization of intestinal microbiota and response to dietary virginiamycin supplementation in the broiler chicken. Appl Environ Microbiol 72:2815–2823.

Dumonceaux TJ, Links MG, Town JR, Hill JE, and Hemmingsen SM. 2017. Targeted capture of *cpn60* gene fragments for PCR-independent microbial community profiling. Protoc exch: Nature publishing group.

Dumonceaux TJ, Schellenberg J, Goleski V, Hill JE, Jaoko W, Kimani J, Money D, Ball TB, Plummer FA, and Severini A. 2009. Multiplex detection of bacteria associated with normal microbiota and with bacterial vaginosis in vaginal swabs by use of oligonucleotide-coupled fluorescent microspheres. J Clin Microbiol 47:4067–4077. 10.1128/jcm.00112-09

Edgar RC. 2017. Accuracy of microbial community diversity estimated by closed- and open-reference OTUs. PeerJ 5:e3889. 10.7717/peerj.3889

Edge TA, Baird DJ, Bilodeau G, Gagné N, Greer C, Konkin D, Newton G, Séguin A, Beaudette L, Bilkhu S, Bush A, Chen W, Comte J, Condie J, Crevecoeur S, El-Kayssi N, Emilson EJS, Fancy D-L, Kandalaft I, Khan IUH, King I, Kreutzweiser D, Lapen D, Lawrence J, Lowe C, Lung O, Martineau C, Meier M, Ogden N, Paré D, Phillips L, Porter TM, Sachs J, Staley Z, Steeves R, Venier L, Veres T, Watson C, Watson S, and Macklin J. 2020. The Ecobiomics project: Advancing metagenomics assessment of soil health and freshwater quality in Canada. Science of The Total Environment 710:135906. https://doi.org/10.1016/j.scitotenv.2019.135906

Fierer N. 2017. Embracing the unknown: disentangling the complexities of the soil microbiome. Nature reviews Microbiology.

Fierer N, Leff JW, Adams BJ, Nielsen UN, Bates ST, Lauber CL, Owens S, Gilbert JA, Wall DH, and Caporaso JG. 2012. Cross-biome metagenomic analyses of soil microbial communities and their functional attributes. Proc Natl Acad Sci USA 109:21390–21395. 10.1073/pnas.1215210110

Gasc C, and Peyret P. 2018. Hybridization capture reveals microbial diversity missed using current profiling methods. Microbiome 6:61. 10.1186/s40168-018-0442-3

Gopal M, and Gupta A. 2016. Microbiome selection could spur next-generation plant breeding strategies. Frontiers in microbiology 7. 10.3389/fmicb.2016.01971

Green SJ, Venkatramanan R, and Naqib A. 2015. Deconstructing the polymerase chain reaction: Understanding and correcting bias associated with primer degeneracies and primer-template mismatches. PLoS ONE 10. 10.1371/journal.pone.0128122

Guo J, Cole JR, Zhang Q, Brown CT, and Tiedje JM. 2016. Microbial community analysis with ribosomal gene fragments from shotgun metagenomes. Appl Environ Microbiol 82:157–166. 10.1128/aem.02772-15

Handley KM, Verberkmoes NC, Steefel CI, Williams KH, Sharon I, Miller CS, Frischkorn KR, Chourey K, Thomas BC, Shah MB, Long PE, Hettich RL, and Banfield JF. 2013. Biostimulation induces syntrophic interactions that impact C, S and N cycling in a sediment microbial community. ISME J 7:800–816. 10.1038/ismej.2012.148

Hebert PDN, Cywinska A, Ball SL, and deWaard JR. 2003. Biological identifications through DNA barcodes. Proc R Soc Lond [Biol] 270:313–321. 10.1098/rspb.2002.2218

Hess M, Sczyrba A, Egan R, Kim TW, Chokhawala H, Schroth G, Luo S, Clark DS, Chen F, Zhang T, Mackie RI, Pennacchio LA, Tringe SG, Visel A, Woyke T, Wang Z, and Rubin EM. 2011. Metagenomic discovery of biomass-degrading genes and genomes from cow rumen. Science 331:463–467.

Hill JE, Hemmingsen SM, Goldade BG, Dumonceaux TJ, Klassen J, Zijlstra RT, Goh SH, and Van Kessel AG. 2005. Comparison of ileum microflora of pigs fed corn-, wheat-, or barley-based diets by chaperonin-60 sequencing and quantitative PCR. Appl Environ Microbiol 71:867–875.

Hill JE, Penny SL, Crowell KG, Goh SH, and Hemmingsen SM. 2004. cpnDB: A chaperonin sequence database. Genome Res 14:1669–1675.

Hill JE, Town JR, and Hemmingsen SM. 2006. Improved template representation in *cpn60* polymerase chain reaction (PCR) product libraries generated from complex templates by application of a specific mixture of PCR primers. Environ Microbiol 8:741–746. 10.1111/j.1462-2920.2005.00944.x

Hill JE, and Vancuren SJ. 2019. Update on cpnDB: a reference database of chaperonin sequences. Database 2019. 10.1093/database/baz033

J T Staley a, and Konopka A. 1985. Measurement of *in situ* activities of nonphotosynthetic microorganisms in aquatic and terrestrial habitats. Annu Rev Microbiol 39:321–346. 10.1146/annurev.mi.39.100185.001541

Johnson LA, Chaban B, Harding JC, and Hill JE. 2015. Optimizing a PCR protocol for cpn60-based microbiome profiling of samples variously contaminated with host genomic DNA. BMC research notes 8:253. 10.1186/s13104-015-1170-4

Kau AL, Ahern PP, Griffin NW, Goodman AL, and Gordon JI. 2011. Human nutrition, the gut microbiome and the immune system. Nature 474:327–336. 10.1038/nature10213

Knight R, Vrbanac A, Taylor BC, Aksenov A, Callewaert C, Debelius J, Gonzalez A, Kosciolek T, McCall L-I, McDonald D, Melnik AV, Morton JT, Navas J, Quinn RA, Sanders JG, Swafford AD, Thompson LR, Tripathi A, Xu ZZ, Zaneveld JR, Zhu Q, Caporaso JG, and Dorrestein PC. 2018. Best practices for analysing microbiomes. Nat Rev Microbiol 16:410–422. 10.1038/s41579-018-0029-9

Koch C, Müller S, Harms H, and Harnisch F. 2014. Microbiomes in bioenergy production: From analysis to management. Curr Opin Biotechnol 27:65–72. 10.1016/j.copbio.2013.11.006

Langmead B, Trapnell C, Pop M, and Salzberg SL. 2009. Ultrafast and memory-efficient alignment of short DNA sequences to the human genome. Genome Biol 10:R25.

Lee CK, Herbold CW, Polson SW, Wommack KE, Williamson SJ, McDonald IR, and Cary SC. 2012. Groundtruthing next-gen sequencing for microbial ecology-biases and errors in community structure estimates from PCR amplicon pyrosequencing. PLoS ONE 7. 10.1371/journal.pone.0044224

Lee DH, Zo YG, and Kim SJ. 1996. Nonradioactive method to study genetic profiles of natural bacterial communities by PCR-single-strand-conformation polymorphism. Appl Environ Microbiol 62:3112–3120.

Links MG, Chaban B, Hemmingsen S, Muirhead K, and Hill J. 2013. mPUMA: a computational approach to microbiota analysis by de novo assembly of operational taxonomic units based on protein-coding barcode sequences. Microbiome 1:23.

Links MG, Demeke T, Gräfenhan T, Hill JE, Hemmingsen SM, and Dumonceaux TJ. 2014. Simultaneous profiling of seed-associated bacteria and fungi reveals antagonistic interactions between microorganisms within a shared epiphytic microbiome on *Triticum* and *Brassica* seeds. New Phytol 202:542–553. 10.1111/nph.12693

Links MG, Dumonceaux TJ, Hemmingsen SM, and Hill JE. 2012. The chaperonin-60 universal target is a barcode for bacteria that enables *de novo* assembly of metagenomic sequence data. PLoS One 7:e49755. 10.1371/journal.pone.0049755

Luo C, Rodriguez-R LM, Johnston ER, Wu L, Cheng L, Xue K, Tu Q, Deng Y, He Z, Shi JZ, Yuan MM, Sherry RA, Li D, Luo Y, Schuur EAG, Chain P, Tiedje JM, Zhou J, and Konstantinidis KT. 2014. Soil microbial community responses to a decade of warming as revealed by comparative metagenomics. Appl Environ Microbiol 80:1777–1786. 10.1128/aem.03712-13

Lynch MDJ, and Neufeld JD. 2015. Ecology and exploration of the rare biosphere. Nat Rev Micro 13:217–229. 10.1038/nrmicro3400

Magoc T, and Salzberg SL. 2011. FLASH: fast length adjustment of short reads to improve genome assemblies. Bioinformatics 27:2957–2963. 10.1093/bioinformatics/btr507

Martin-Laurent F, Marti R, Waglechner N, Wright GD, and Topp E. 2014. Draft genome sequence of the sulfonamide antibiotic-degrading *Microbacterium* sp. strain C448. Genome Announc 2. 10.1128/genomeA.01113-13

Martin M. 2011. Cutadapt removes adapter sequences from high-throughput sequencing reads. 2011 17:3. 10.14806/ej.17.1.200

McKenney EA, Ashwell M, Lambert JE, and Fellner V. 2014. Fecal microbial diversity and putative function in captive western lowland gorillas (*Gorilla gorilla* gorilla), common chimpanzees (*Pan troglodytes*), Hamadryas baboons (*Papio hamadryas*) and binturongs (*Arctictis binturong*). Int Zool 9:557–569. 10.1111/1749-4877.12112

Muegge BD, Kuczynski J, Knights D, Clemente JC, González A, Fontana L, Henrissat B, Knight R, and Gordon JI. 2011. Diet drives convergence in gut microbiome functions across mammalian phylogeny and within humans. Science 332:970–974. 10.1126/science.1198719

Pérez-López E, Hammond C, Olivier CY, and Dumonceaux TJ. 2017. Detection and Typing of ‘Candidatus Phytoplasma’ spp. in Host DNA extracts Using Oligonucleotide-Coupled Fluorescent Microspheres. In: Bishop-Lilly KA, ed. Diagnostic Bacteriology. New York: Humana Press.

Peterson SW, Knox NC, Golding GR, Tyler SD, Tyler AD, Mabon P, Embree JE, Fleming F, Fanella S, Van Domselaar G, Mulvey MR, and Graham MR. 2016. A Study of the Infant Nasal Microbiome Development over the First Year of Life and in Relation to Their Primary Adult Caregivers Using *cpn60* Universal Target (UT) as a Phylogenetic Marker. PLoS ONE 11:e0152493. 10.1371/journal.pone.0152493

Pinto AJ, and Raskin L. 2012. PCR biases distort bacterial and archaeal community structure in pyrosequencing datasets. PLOS ONE 7:e43093. 10.1371/journal.pone.0043093

Poretsky R, Rodriguez-R LM, Luo C, Tsementzi D, and Konstantinidis KT. 2014. Strengths and limitations of 16S rRNA gene amplicon sequencing in revealing temporal microbial community dynamics. PLOS ONE 9:e93827. 10.1371/journal.pone.0093827

Props R, Kerckhof F-M, Rubbens P, De Vrieze J, Hernandez Sanabria E, Waegeman W, Monsieurs P, Hammes F, and Boon N. 2016. Absolute quantification of microbial taxon abundances. ISME J. 10.1038/ismej.2016.117

Ranjan R, Rani A, Metwally A, McGee HS, and Perkins DL. 2016. Analysis of the microbiome: Advantages of whole genome shotgun versus 16S amplicon sequencing. Biochem Biophys Res Commun 469:967–977. http://dx.doi.org/10.1016/j.bbrc.2015.12.083

Raymond F, Ouameur AA, Deraspe M, Iqbal N, Gingras H, Dridi B, Leprohon P, Plante P-L, Giroux R, Berube E, Frenette J, Boudreau DK, Simard J-L, Chabot I, Domingo M-C, Trottier S, Boissinot M, Huletsky A, Roy PH, Ouellette M, Bergeron MG, and Corbeil J. 2016. The initial state of the human gut microbiome determines its reshaping by antibiotics. ISME J 10:707–720. 10.1038/ismej.2015.148

Robertson G, Schein J, Chiu R, Corbett R, Field M, Jackman SD, Mungall K, Lee S, Okada HM, Qian JQ, Griffith M, Raymond A, Thiessen N, Cezard T, Butterfield YS, Newsome R, Chan SK, She R, Varhol R, Kamoh B, Prabhu A-L, Tam A, Zhao Y, Moore RA, Hirst M, Marra MA, Jones SJM, Hoodless PA, and Birol I. 2010. De novo assembly and analysis of RNA-seq data. Nature Meth 7:909–912. 10.1038/nmeth.1517

Rozen S, and Skaletsky H. 2000. Primer3 on the WWW for general users and for biologist programmers. Methods in molecular biology (Clifton, NJ) 132:365–386.

Schoch CL, Seifert KA, Huhndorf S, Robert V, Spouge JL, Levesque CA, Chen W, and Fungal Barcoding C. 2012. Nuclear ribosomal internal transcribed spacer (ITS) region as a universal DNA barcode marker for Fungi. Proc Natl Acad Sci USA 109:6241–6246. 10.1073/pnas.1117018109

Schuenemann VJ, Bos K, DeWitte S, Schmedes S, Jamieson J, Mittnik A, Forrest S, Coombes BK, Wood JW, Earn DJD, White W, Krause J, and Poinar HN. 2011. Targeted enrichment of ancient pathogens yielding the pPCP1 plasmid of *Yersinia pestis* from victims of the Black Death. Proc Natl Acad Sci USA 108:E746–E752. 10.1073/pnas.1105107108

Schwartz MH, Wang H, Pan JN, Clark WC, Cui S, Eckwahl MJ, Pan DW, Parisien M, Owens SM, Cheng BL, Martinez K, Xu J, Chang EB, Pan T, and Eren AM. 2018. Microbiome characterization by high-throughput transfer RNA sequencing and modification analysis. Nature Communications 9:5353. 10.1038/s41467-018-07675-z

Singer E, Bushnell B, Coleman-Derr D, Bowman B, Bowers RM, Levy A, Gies EA, Cheng J-F, Copeland A, Klenk H-P, Hallam SJ, Hugenholtz P, Tringe SG, and Woyke T. 2016. High-resolution phylogenetic microbial community profiling. ISME J 10:2020–2032. 10.1038/ismej.2015.249

Tikhonovich IA, and Provorov NA. 2011. Microbiology is the basis of sustainable agriculture: An opinion. Ann Appl Biol 159:155–168. 10.1111/j.1744-7348.2011.00489.x

Topp E, Chapman R, Devers-Lamrani M, Hartmann A, Marti R, Martin-Laurent F, Sabourin L, Scott A, and Sumarah M. 2013. Accelerated biodegradation of veterinary antibiotics in agricultural soil following long-term exposure, and isolation of a sulfamethazine-degrading *Microbacterium* sp. J Environ Qual 42:173–178. 10.2134/jeq2012.0162

Town JR, Links MG, Fonstad TA, and Dumonceaux TJ. 2014. Molecular characterization of anaerobic digester microbial communities identifies microorganisms that correlate to reactor performance. Bioresour Technol 151:249–257. http://dx.doi.org/10.1016/j.biortech.2013.10.070

Turnbaugh PJ, Ley RE, Hamady M, Fraser-Liggett CM, Knight R, and Gordon JI. 2007. The Human Microbiome Project. Nature 449:804–810. 10.1038/nature06244

Vancuren SJ, Dos Santos SJ, Hill JE, and the Maternal Microbiome Legacy Project T. 2020. Evaluation of variant calling for cpn60 barcode sequence-based microbiome profiling. PLOS ONE 15:e0235682. 10.1371/journal.pone.0235682

Wagner DM, Klunk J, Harbeck M, Devault A, Waglechner N, Sahl JW, Enk J, Birdsell DN, Kuch M, Lumibao C, Poinar D, Pearson T, Fourment M, Golding B, Riehm JM, Earn DJD, DeWitte S, Rouillard J-M, Grupe G, Wiechmann I, Bliska JB, Keim PS, Scholz HC, Holmes EC, and Poinar H. 2014. *Yersinia pestis* and the Plague of Justinian 541-543 AD: a genomic analysis. Lancet Infect Dis.

Walker AW, Martin JC, Scott P, Parkhill J, Flint HJ, and Scott KP. 2015. 16S rRNA gene-based profiling of the human infant gut microbiota is strongly influenced by sample processing and PCR primer choice. Microbiome 3:26. 10.1186/s40168-015-0087-4

Weller R, and Ward DM. 1989. Selective recovery of 16S rRNA sequences from natural microbial communities in the form of cDNA. Appl Environ Microbiol 55:1818–1822.

Woese CR, and Fox GE. 1977. Phylogenetic structure of the prokaryotic domain: The primary kingdoms. Proc Natl Acad Sci USA 74:5088–5090. 10.1073/pnas.74.11.5088

Woese CR, Kandler O, and Wheelis ML. 1990. Towards a natural system of organisms: proposal for the domains Archaea, Bacteria, and Eucarya. Proc Natl Acad Sci USA 87:4576–4579. 10.1073/pnas.87.12.4576

