## Supplementary figures and images for "CaptureSeq: Hybridization-based enrichment of *cpn60* gene fragments reveals the community structures of synthetic and natural microbial ecosystems"

### Supplemental Fig S1

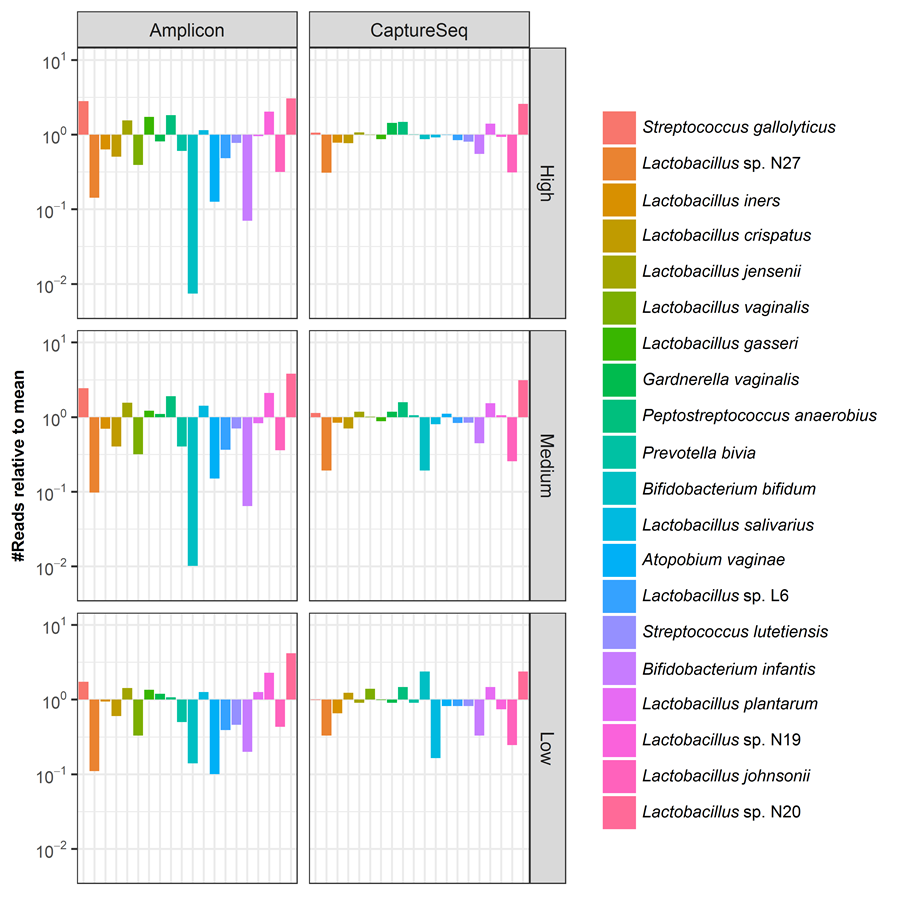
