## Supplementary material for "CaptureSeq: Hybridization-based enrichment of *cpn60* gene fragments reveals the community structures of synthetic and natural microbial ecosystems": Table S2

|  | High | Medium | Low |
| --- | --- | --- | --- |
| *Atopobium vaginae* | 943 | 87 | 10 |
| *Bifidobacterium bifidum* | 827 | 15 | 29 |
| *Bifidobacterium infantis* | 527 | 35 | 4 |
| *Gardnerella vaginalis* | 1,369 | 92 | 11 |
| *Lactobacillus* sp. N27 | 295 | 15 | 4 |
| *Lactobacillus iners* | 741 | 66 | 8 |
| *Lactobacillus crispatus* | 731 | 55 | 15 |
| *Lactobacillus jensenii* | 1,020 | 92 | 11 |
| *Lactobacillus vaginalis* | 946 | 79 | 17 |
| *Lactobacillus gasseri* | 827 | 69 | 12 |
| *Lactobacillus salivarius* | 874 | 63 | 2 |
| *Lactobacillus* sp. L6 | 803 | 65 | 10 |
| *Lactobacillus* sp. N19 | 889 | 83 | 9 |
| *Lactobacillus johnsonii* | 297 | 20 | 3 |
| *Lactobacillus* sp. N20 | 2,466 | 244 | 29 |
| *Peptostreptococcus anaerobius* | 1,416 | 124 | 18 |
| *Prevotella bivia* | 959 | 83 | 11 |
| *Streptococcus gallolyticus* | 1,010 | 89 | 12 |
| *Streptococcus lutetiensis* | 769 | 66 | 10 |
| *Lactobacillus plantarum* | 1,332 | 120 | 18 |

**Table S2.** Sequencing read abundances mapping to each taxonomic cluster for seed wash samples spiked with a synthetic community consisting of 20 *cpn60* UT plasmids in 10-fold decreasing dilutions (high, medium, low) as described in the text. CaptureSeq data were downsampled to 506,247 sequencing reads and mapped to a reference dataset consisting of the *cpn60* UT sequences of the 20 bacteria in the panel.
