## Supplementary material for "CaptureSeq: Hybridization-based enrichment of *cpn60* gene fragments reveals the community structures of synthetic and natural microbial ecosystems": Table S3

| Profiling method | Treatment  (mg kg^-1^) | Plot | Sequencing  reads | Reference Mapped reads |
| --- | --- | --- | --- | --- |
| CaptureSeq | 0 | 1 | 1,226,019 | 16.4% |
|  | 0 | 7 | 1,172,125 | 18.1% |
|  | 0 | 12 | 1,158,656 | 15.9% |
|  | 10 | 4 | 1,025,680 | 17.0% |
|  | 10 | 8 | 827,129 | 16.4% |
|  | 10 | 11 | 765,101 | 16.7% |
| Shotgun metagenomic | 0 | 1 | 6,116,268 | 0.070% |
|  | 0 | 7 | 7,306,386 | 0.071% |
|  | 0 | 12 | 6,953,220 | 0.070% |
|  | 10 | 4 | 4,232,651 | 0.067% |
|  | 10 | 8 | 4,794,243 | 0.067% |
|  | 10 | 11 | 4,287,559 | 0.069% |

**Table S2.** Read numbers (total and proportion mapping to the *cpn60* reference dataset) obtained using CaptureSeq and shotgun metagenomic profiling methods on soil samples.
